# Allele frequency dynamics in a pedigreed natural population

**DOI:** 10.1101/388710

**Authors:** Nancy Chen, Ivan Juric, Elissa J. Cosgrove, Reed Bowman, John W. Fitzpatrick, Stephan J. Schoech, Andrew G. Clark, Graham Coop

## Abstract

A central goal of population genetics is to understand how genetic drift, natural selection, and gene flow shape allele frequencies through time. However, the actual processes underlying these changes - variation in individual survival, reproductive success, and movement - are often difficult to quantify. Fully understanding these processes requires the population pedigree, the set of relationships among all individuals in the population through time. Here, we use extensive pedigree and genomic information from a long-studied natural population of Florida Scrub-Jays (*Aphelocoma coerulescens*) to directly characterize the relative roles of different evolutionary processes in shaping patterns of genetic variation through time. We performed gene dropping simulations to estimate individual genetic contributions to the population and model drift on the known pedigree. We found that observed allele frequency changes are generally well predicted by accounting for the different genetic contributions of founders. Our results show that the genetic contribution of recent immigrants is substantial, with some large allele frequency shifts that otherwise may have been attributed to selection actually due to gene flow. We identified a few SNPs under directional short-term selection after appropriately accounting for gene flow. Using models that account for changes in population size, we partitioned the proportion of variance in allele frequency change through time. Observed allele frequency changes are primarily due to variation in survival and reproductive success, with gene flow making a smaller contribution. This study provides one of the most complete descriptions of short-term evolutionary change in allele frequencies in a natural population to date.

---

An evolving natural population is essentially a vast pedigree, with genetic material transmitted down this pedigree following the laws of Mendelian inheritance (except in rare cases of meiotic drive). We often cannot directly observe the actual processes underlying genetic change. Instead, population genetic studies typically rely on current day patterns of genetic variation - or, if temporal samples are available, the variation in allele frequencies through time - to make inferences about the effects of genetic drift, natural selection, and gene flow in driving evolutionary change. However, these evolutionary mechanisms can be precisely understood in terms of the differential genetic contributions of individuals to the population pedigree over time, combined with the stochasticity of Mendelian segregation.

Knowledge of the population pedigree allows us to trace expected individual genetic contributions, *i.e*., the expected number of copies of a neutral allele contributed by a given individual, to the population in future generations. Individual genetic contributions can be estimated analytically (1–3) or via gene dropping simulations, i.e., simulations of Mendelian transmission of alleles down their pedigree of descendants (4). The long-term genetic contribution of an individual is an individual’s reproductive value, a general measure of individual fitness (5–7). Indeed, the reproductive value of an individual influences many aspects of the survival of an individual’s genotype, from the probability of loss of a new, weakly beneficial mutation to the complex distribution of genomic blocks passed on to future generations (8).

Analyses of known pedigrees have been used to estimate individual genetic contributions to assess founder effects in human populations (1–3, 9–11) and to predict the probability of gene loss in captive breeding populations (4, 12). Also, empirical pedigree calculations have long been used to understand genetic models of human diseases (13) and are increasingly used in natural populations to understand the genetic basis of quantitative trait variation, fitness consequences of inbreeding, and much more (14). To date, empirical pedigrees and gene dropping approaches have been rarely used to study the temporal spread and loss of individual alleles (15–18).

Here, we combine genomic data with a known population pedigree to describe and predict allele frequency change at many loci in an exhaustively sampled free-living population of Florida Scrub-Jays (*Aphelocoma coerulescens*) at Archbold Biological Station. Intensive study since 1969 has resulted in lifetime fitness measures for thousands of individuals on an extensive pedigree. Recently, Chen *et al*. (19) generated genome-wide single nucleotide polymorphism (SNP) data for nearly every individual in the population over the past two decades. In this study, we link individual lifetime reproductive success with long-term genetic contributions and allele frequency change. We show how the population pedigree presents a powerful opportunity to directly elucidate the relative roles of drift, gene flow, and selection in governing allele frequency dynamics over time.

## Results

### Individual fitness and genetic contributions

First, we consider a series of inferences that can be made purely with the pedigree, ignoring the SNP genotypes for the moment. We measured fitness for all 926 individuals who bred in our study population in 1990-2013 and were born before 2002 (the age cohorts who are all dead by the end of 2014). Lifetime reproductive success was highly variable in our study population: the total number of nestlings produced over an individual’s lifetime ranged from 0 to 43, with 197 individuals (21%) producing no nestlings despite having at least one breeding attempt (Fig. S1). Only 43% produced any grandchildren (range 0-189), and 33% produced great-grandchildren (range 0-210). As might be expected in these monogamous birds in which the sexes experience equal annual mortality (20, 21), we found no significant differences in individual fitness between males and females (Wilcoxon rank-sum test, *p* > 0.65 for all three measures of fitness).

Using the detailed population pedigree, we calculated both the genealogical and expected genetic contribution of each individual to the study population from 1990-2013. Fig. 1AB shows results for two illustrative males, both of whom first bred in 1994. Male A lived until 2006 and had 41 offspring, whereas Male B only lived until 2000 and had 7 offspring. We define an individual’s <u>genealogical contribution</u> to a given year as the proportion of nestlings in the birth cohort who are genealogically descended from the focal individual, while an individual’s <u>expected genetic contribution</u> is the expected proportion of alleles at a locus in the nestling cohort that come from the focal individual. Beyond a few generations, few genealogical descendants are expected to inherit genetic material, so the number of genealogical descendants should quickly outnumber the number of genetic descendants. Fig. 1ABC nicely demonstrates this pattern in our data, providing empirical illustration for a substantial body of theory on the relationship between genetic and genealogical ancestry (8, 22, 23). An individual’s genealogical contribution in 2013 is correlated with its expected genetic contribution in 2013 (Spearman’s *ρ* = 0.99, *p* < 2 × 10^−16^), but its genealogical contribution is significantly larger (paired Wilcoxon test, *p* < 2 × 10^−16^; Fig. 1C).

**Figure 1.**
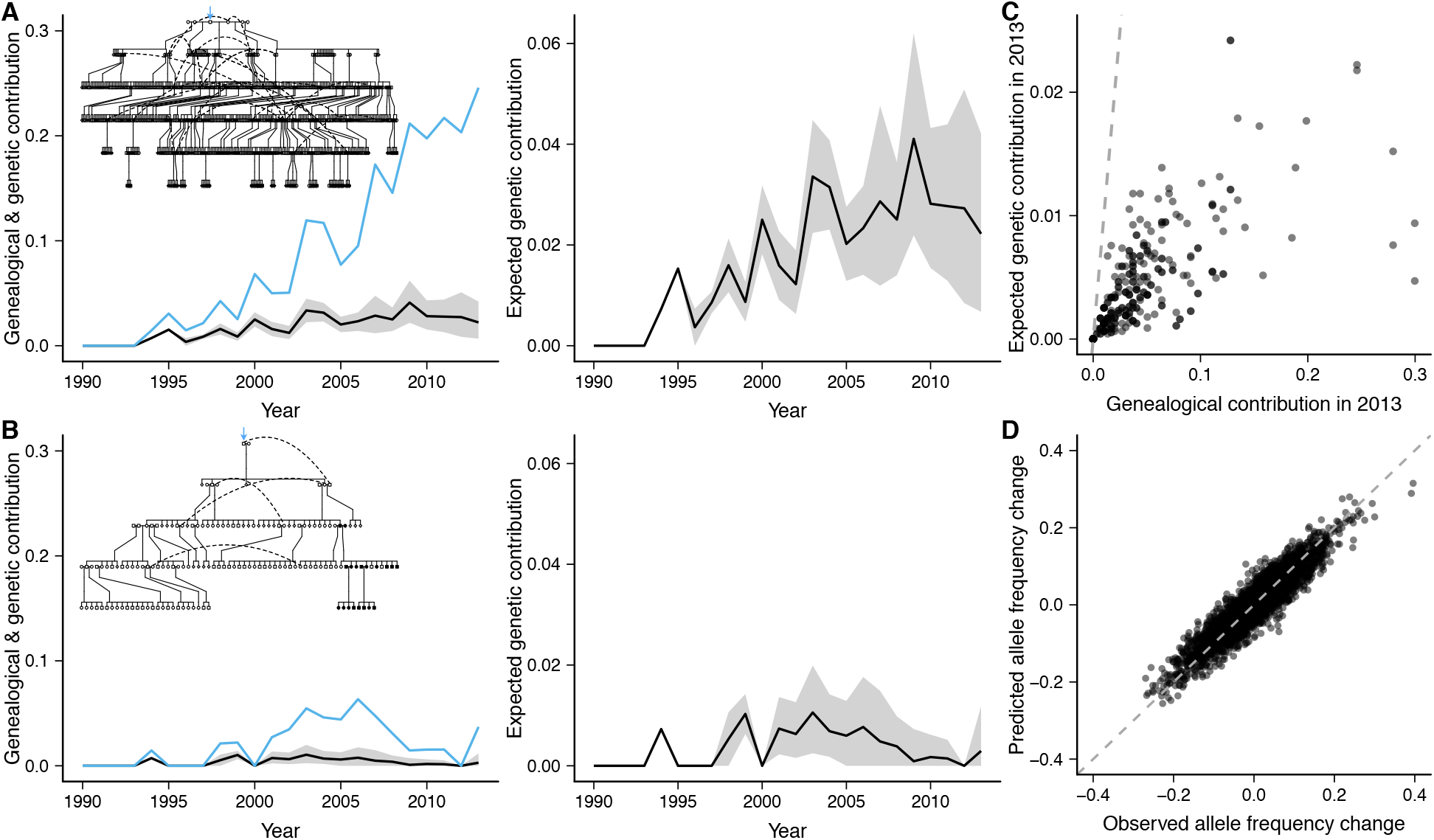
Genealogical and expected genetic contributions to the study population over time for two males who first bred in 1994 with total lifetime reproductive success of (A) 41 and (B) 7. Blue lines indicate the proportion of nestlings each year who are genealogical descendants. Black lines indicate mean expected genetic contribution for each year and grey shading the 95% confidence interval for their contribution at a neutral locus. The pedigree of all descendants of each individual in the study population is shown, with an arrow indicating the focal individual and solid symbols denoting individuals still alive in 2013. (C) Genealogical contributions and expected genetic contributions to the population in 2013 for all breeders born before 2002 and who first bred in 1990 or later (926 individuals). The dotted line indicates a one-to-one relationship. (D) Predicted versus observed change in allele frequencies from 1999 to 2013.

Individual fitness is a central concept in evolutionary biology but notoriously difficult to measure (24). Here, we tested for a relationship between various proxies for fitness and the expected genetic contribution to the population. All three measures of fitness (number of offspring, grandoffspring, and great-grandoffspring) are significantly correlated with both the total expected genetic contribution from 1990-2013 (Spearman’s *ρ* = 0.92, 0.85, 0.78, respectively; *p* < 2 × 10^−16^ for each comparison) and the expected genetic contribution to the 2013 nestling cohort (Spearman’s *ρ* = 0.57, 0.83, 0.87, respectively; *p* < 2 × 10^−16^ for each comparison; Fig. S2). The correlation between individual fitness and expected genetic contribution in 2013 increases with the number of generations considered in the measure of fitness.

### Allele frequency predictions

In previous work, we genotyped >80% of all adults and nearly every nestling born in 1989-1991,1995, and 1999-2013 at 10,731 autosomal SNPs (19). Here, we investigate allele frequency dynamics in the birth cohort from 1999-2013. In theory, we should be able to predict the allele frequency of a particular SNP in a given year simply by summing the individual genetic contributions of each founder to the population that year weighted by the founder’s genotype at that SNP. Note that immigrants are considered founders, so this approach incorporates gene flow. We generated allele frequency predictions for each autosomal SNP in 1999-2013. We can nearly perfectly predict the allele frequency for each SNP in any given year (*β* = 0.99). More importantly, we can predict the overall net change in allele frequencies from 1999 to 2013 (*β* = 0.87; Fig. 1D).

### Effect of gene flow

Previous work showed high levels of immigration into our study population (19), with immigrants comprising 32-55% of all breeding adults in a given year (19). We estimated the cumulative expected genetic contribution of new immigrants appearing in our study population from 1991 onward (Fig. 2A). Total expected genetic contributions of individual immigrant cohorts in 2013 range from 0.003-0.083 and are significantly correlated with the number of individuals in that cohort (Spearman’s *ρ* = 0.52, *p* = 0.01). Immigrants arriving since 1990 are, in aggregate, expected to contribute 75% of the alleles present in the 2013 nestling cohort. We fitted a model to project the contributions of immigrants into the future (Fig. S3). We predict that it takes on average 32 years for 95% of neutral alleles to be replaced by immigration.

**Figure 2.**
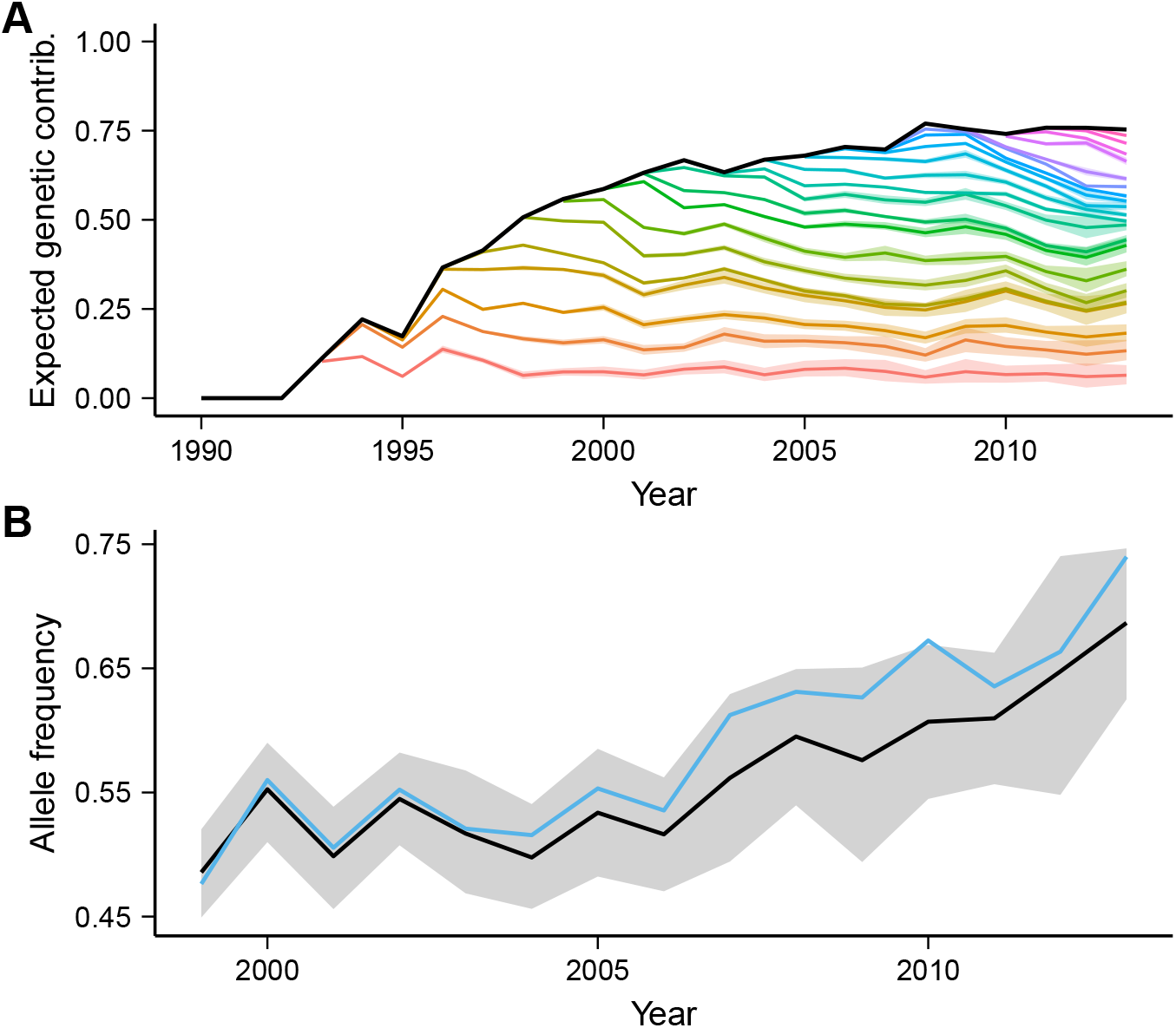
(A) The expected genetic contribution of different cohorts of recent immigrants (based on the year they were first observed in our population). The black line shows the total expected genetic contribution of immigrants appearing in the population after 1990. Each colored line shows the mean added contribution of successive cohorts of immigrants, with shading to show the 95% confidence intervals. (B) Observed (blue) and simulated (black) allele frequencies over time for a SNP with significantly increasing immigrant allele frequencies.

With the high expected genetic contribution of immigrants, we predicted that gene flow could play an important role in governing allele frequency trajectories over time. While the majority of SNPs show small frequency changes, we do observe a few large allele frequency shifts over this 15 year time period: the difference in allele frequencies between 1999 and 2013 is >0.15 for 129 SNPs and >0.2 for 11 SNPs. We used gene dropping simulations to model the expected allele frequency distributions at each SNP in the nestling cohorts from 1999-2013. Unlike our previous pedigree-based simulations to generate individual genetic contributions, here we began simulations with the observed founder genotypes for each SNP. The mean allele frequency of these gene dropping simulations is equal to the allele frequency predictions generated above.

Indeed, we found that gene flow alone can cause large allele frequency shifts (one example shown in Fig. 2B). This allele increased in frequency by 0.26 between 1999-2013, yet the observed allele frequency trajectory lies well within expectations from our gene dropping simulations. For this SNP, the allele frequency in incoming immigrants significantly increased over time (Mann Kendall test, *p* = 0.002), from 0.51 in the 1990 founders to 0.71 in immigrants appearing in 2013, likely causing the population allele frequency to increase as well. As gene dropping begins with founder genotypes, any change in allele frequency due to incoming immigration is reflected in the simulation results. In the absence of data on the pedigree and the genotypes of immigrants, such trajectories could resemble selection, but our gene-dropping approach shows that these large changes in allele frequencies are actually likely the result of gene flow.

### Short-term selection

Given that our gene dropping simulations accurately account for the effects of both gene flow and drift, we then tested for significant net allele frequency changes from 1999-2013 as well as between all adjacent years during this time period. We compared observed allele frequency shifts to the expected distribution of allele frequency shifts generated from the gene dropping simulations (Fig. 3A). At a false discovery rate (FDR) of 0.25, 18 SNPs showed significant changes in allele frequency between 1999 and 2013 (Table S1, Fig. 3). For allele frequency shifts between adjacent years, we find some hits if we treat each year as an independent test (Table S2, Fig. S4); no SNPs survived multiple testing correction across years. Overall, the gene dropping simulations provide a good fit to observed data (Fig. S5), suggesting allele frequency change in our population during this time period is largely consistent with a neutral model.

**Figure 3.**
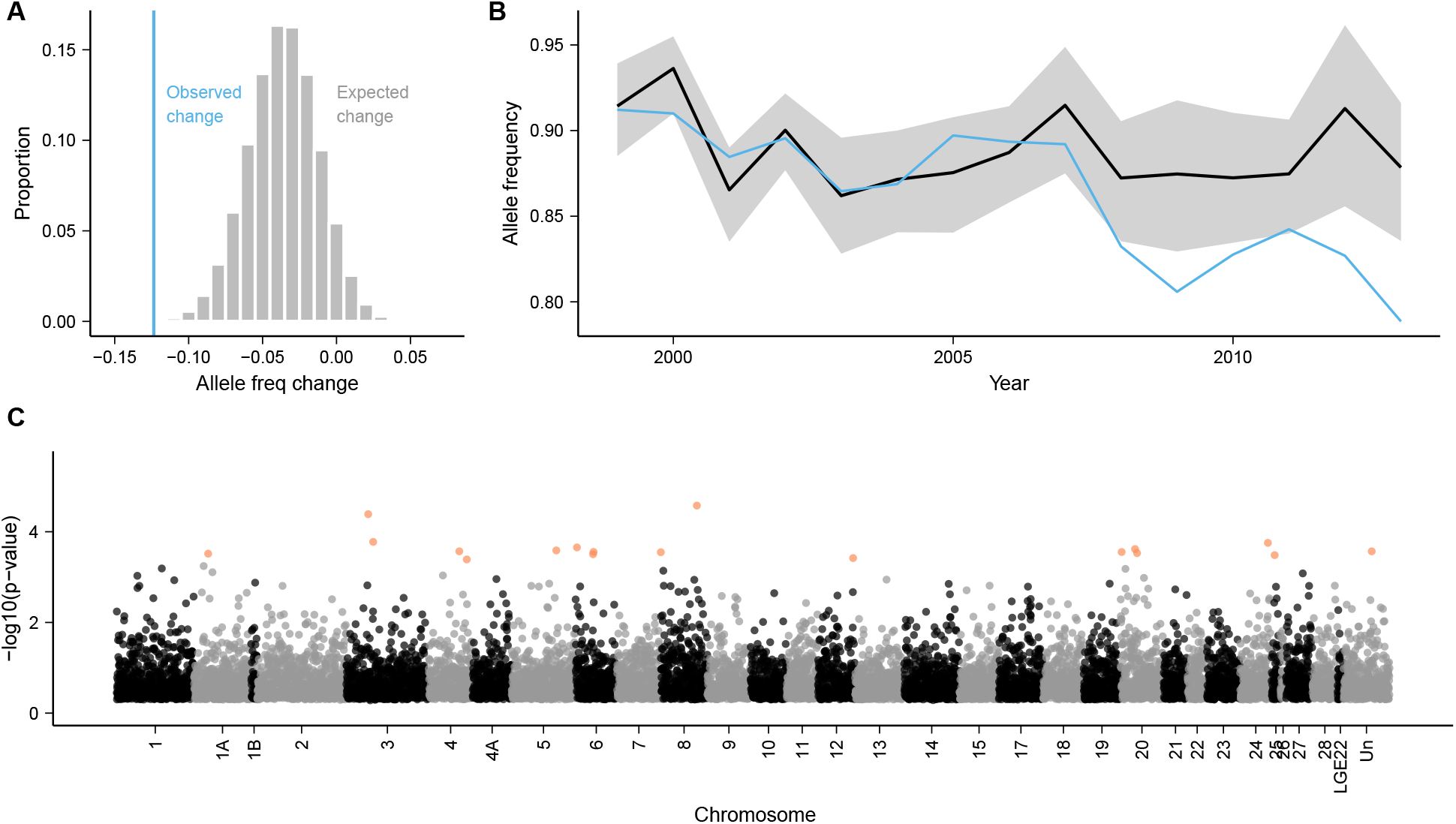
(A) Distribution of expected allele frequency shifts between 1999 and 2013 for the SNP shown in B (grey histogram). The blue line indicates the observed allele frequency change. (B) Observed (blue) and simulated (black) allele frequency trajectories for one of the significant SNPs in 1999-2013. Grey bars indicate 95% confidence intervals for the gene dropping simulations. (C) Manhattan plot for allele frequency shifts in 1999-2013. Significant SNPs (FDR < 0.25) are highlighted in orange.

### Variance in allele frequencies through time

Finally, to quantify the relative roles of different evolutionary processes in shaping patterns of genetic variation genome-wide, we constructed a model for the variance in allele frequency change in 1999-2013. We assume that allele frequencies change due to just three processes: differential survival of individuals, immigration, and reproduction. We partitioned the proportion of allele frequency change from year to year due to survival/reproduction and gene flow using a model that accounts for variation in population sizes over time and overlapping generations (Fig. 4). The change in allele frequency due to births is a result of both variation in family size and Mendelian segregation of alleles in heterozygotes. We further divided the variance in allele frequency change due to births into these two components and found that the noise due to Mendelian segregation comprises 24-48% of the variance due to births, and 12-23% of the overall variance. Our model results reflect patterns we observed in the field. For instance, the number of nestlings born in 2012 was unusually low (Fig. S6), leading the survivors to have a disproportionate impact on allele frequency variation in 2011-2012. Overall, we found that 90% of the variance in allele frequencies is driven by variation in survival and reproductive success among individuals, which is consistent with our small population size.

**Figure 4.**
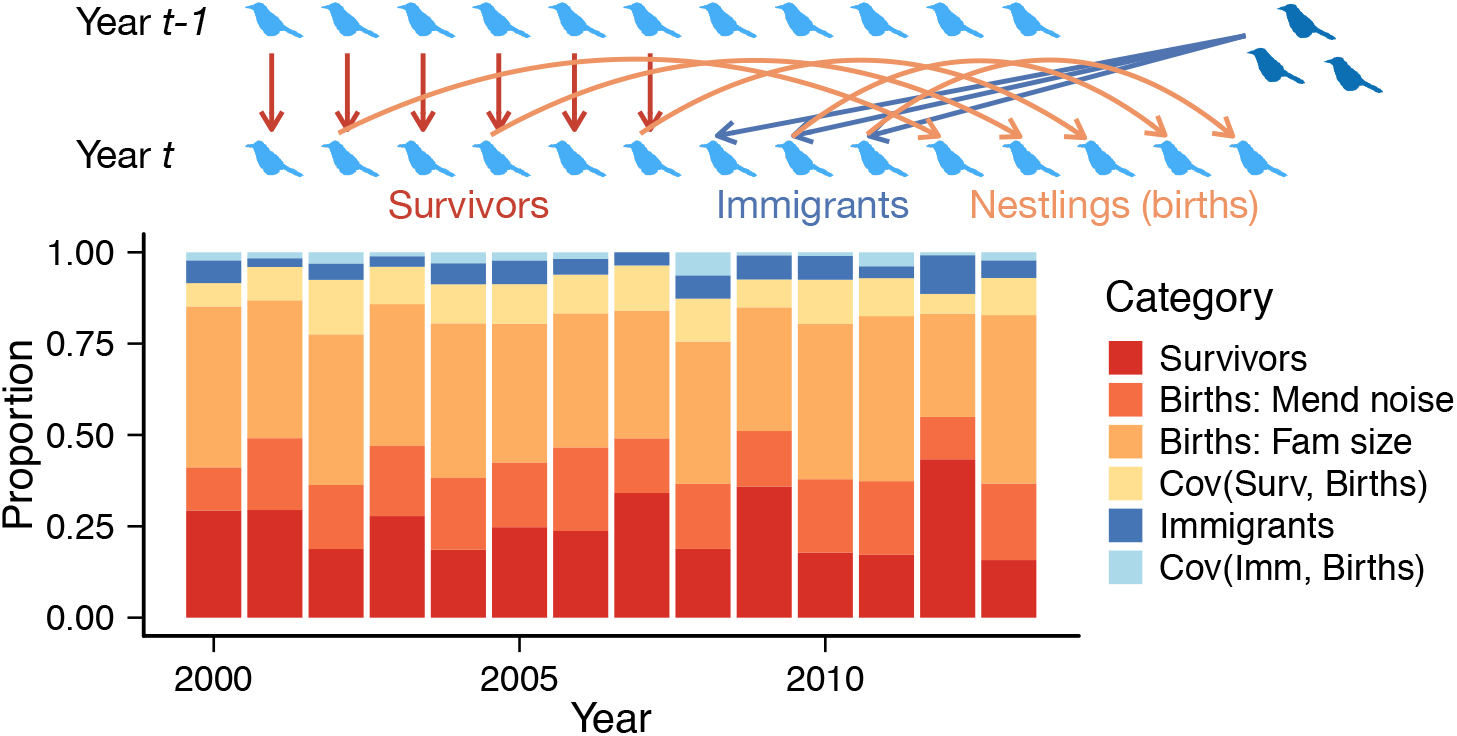
Schematic and results for our model of the variance in autosomal allele frequency change from year to year due to survival/reproduction (red/orange/yellow) or gene flow (blue). The variance in allele frequency change due to births is further partitioned into the variance due to variation in family size and additional noise due to Mendelian segregation of heterozygotes.

## Discussion

We capitalized on a long-term demographic study of Florida Scrub-Jays with extensive pedigree and genomic data to demonstrate how short-term evolutionary processes operate in a natural population. We estimated genealogical and expected genetic contributions for hundreds of individuals, and linked genetic contributions to both individual fitness measures and allele frequency change over time. In our population of Florida Scrub-Jays, we observed huge variation in individual fitness: 75% of the 445 individuals who first bred in our population before 1997 have no living descendants by 2013, but 6 of these individuals are each genealogical ancestors to >25% of the birth cohort in 2013. However, many of these genealogical descendants receive little genetic material from a particular ancestor, thanks to the vagaries of Mendelian segregation and recombination during meiosis (8, 22, 23). Here, we empirically show how genealogical contributions outstrip expected genetic contributions after just a few generations.

Individual fitness is defined as an individual’s genetic contribution to future generations but is typically measured using single-generation proxies such as lifetime reproductive success. Similar to (24), we found that lifetime reproductive success is correlated with an individual’s expected genetic contribution to the population in the future. Florida Scrub-Jays rarely move once they become an established breeder on a territory, giving us confidence in our measures of total lifetime reproductive success. Our estimates of the total number of grand-offspring or great-grandoffspring, however, may be an underestimate because a few of the individuals in our sample still have surviving children and any descendants of emigrants are not counted. We believe the latter is a minor issue because we know that emigration rates are extremely low from annual surveys of the surrounding areas. The correlation between the number of descendants and expected genetic contribution in 2013 is higher for fitness proxies that include more generations. Longer-term fitness proxies can be more accurate in part because they include variation in offspring quality (24), an idea we could explore by estimating genetic correlation of number of offspring and the number of grandoffspring (25).

The high expected genetic contribution of immigrants is consistent with previous results showing that immigrants play an important role in maintaining levels of genetic variation in the population (19). Genome-wide, allele frequency changes are primarily driven by variation in individual survival and reproduction. The contribution of new immigrants to allele frequency changes from year to year (Fig. 4) and is much smaller than the cumulative expected genetic contribution of immigrants compounded over generations (Fig. 2A). This discrepancy occurs because in our model, immigrants are included in allele frequency change only in the year they appear, while their genetic contributions to future years is folded into variation in survival and births. The change in allele frequencies we see due to variation in survival and births, except for the deviation due to Mendelian segregation of heterozygotes, includes the contribution of natural selection, and so these proportions should be viewed as including the contributions of both drift and selection to allele frequency change.

We used gene dropping to predict allele frequency changes over time for individual SNPs across the genome and showed that SNP trajectories can sometimes be strongly driven by gene flow. Our results emphasize the importance of knowing the underlying demography of population, as large allele frequency shifts that ordinarily may be attributed to selection could be due to processes such as drift and gene flow. Though we did detect signatures of selection changing allele frequencies in a few adjacent years, overall we found little evidence of strong directional selection on single alleles on this short timescale.

One of the reasons why we detect so few selected loci is the accuracy with which we can predict allele frequency change from individual genetic contributions and observed founder genotypes. By conditioning on the population pedigree and founder genotypes, our gene-dropping simulations appropriately accounted for variation in population sizes over time and relatedness within the birth cohort, as well as the effects of gene flow. One could argue that using gene dropping to test for selection is conservative, as the pedigree itself encodes information about variation in fitness. However, variation in offspring number is a natural part of genetic drift (26), while heritable variation in fitness at unlinked loci can act to compound genetic drift over the generations (27–29). Therefore, gene-dropping simulations on the population pedigree provide the correct null model for heritable fitness variation for neutral alleles are that are unlinked to selected alleles.

Here we have traced only single alleles down the pedigree. The incorporation of linkage and haplotype information would allow the quantification of realized, actual genetic contributions for each individual instead of just expected genetic contributions. By tracing the inheritance of genomic blocks down the pedigree, we could explore the relationship between reproductive value and the distribution of surviving genetic material, quantify the actual genetic contribution of recent immigrants across the genome, as well as pinpoint specific haplotypes linked to fitness. However, even single SNP analyses on a population pedigree provide substantial insights to the evolutionary forces governing allele frequency dynamics over time. As genomic resources for pedigreed populations expand, our ability to directly observe the causes and consequences of short-term evolution will increase dramatically.

## Materials and Methods

### Study system and dataset

The Florida Scrub-Jay is a non-migratory, cooperatively breeding bird restricted to oak scrub in Florida (30). A population of Florida Scrub-Jays has been intensively monitored at Archbold Biological Station (Venus, Florida, USA) for decades. Woolfenden, Fitzpatrick, Bowman, and colleagues began monitoring the northern half in 1969 (30), and Mumme, Schoech, and colleagues began monitoring the southern half in 1989 (31, 32). All individuals in the entire population are uniquely banded, allowing identification of immigrant individuals each year. The entire population is censused every few months and all nests of all family groups are closely monitored, providing documentation of survival and reproductive success for all individuals in the population. All fieldwork was approved by the Institutional Animal Care and Use Committees at Cornell University (IACUC 2010-0015), the University of Memphis (0667), and Archbold Biological Station (AUP-006-R) and permitted by the U.S. Fish and Wildlife Service (TE824723-8, TE-117769), the US Geological Survey (banding permits 07732, 23098)), and the Florida Fish and Wildlife Conservation Commission (LSSC-10-00205).

Because of the very low rate of extra-pair paternity and limited natal dispersal distances in this population (19–21, 33), we have a detailed and accurate population pedigree. To avoid any artifacts caused by study tract expansion before 1990, we began all our analyses in 1990 and truncated the pedigree accordingly. Our final pedigree consists of 6,936 individuals. We used the pedigree to estimate individual fitness for all adults who first bred in 1990 or later and were born before 2002 (926 individuals) by counting the total number of offspring, grandoffspring, or great-grandoffspring produced by a given individual over its lifetime. We restricted our sample to age cohorts of breeders who all died before the end of 2014 to ensure an accurate and unbiased survey of lifetime reproductive success. Of these individuals, 5% had children who were still alive at the end of 2014 and may produce additional grandchildren, and 13% had grandchildren who were still alive at the end of 2014 and therefore may produce additional great-grandchildren. Here, we define offspring as 11-day-old nestlings (the age at which they are first banded).

For our genomic analyses, we focused on a core set of approximately 68 territories in a geographic area that has been consistently monitored starting in 1990. In a previous study, we genotyped 3,984 individuals at 15,416 genome-wide SNPs, resulting in near-complete sampling of all nestlings and breeders in these core territories in 1989-1991, 1995, and 1999-2013 (19). Information on SNP discovery, genotyping, and pedigree verification can be found in (19). Here, we removed SNPs with minor allele frequency < 0.05. Our final dataset consists of 10,731 autosomal SNPs in 3,404 individuals. All data used in this study can be found at Figshare.

### Expected genetic contributions

We quantify individual genetic contribution as the expected proportion of alleles in the nestling cohort that comes from the focal individual. The expected genetic contribution of an individual to a given year can be calculated as:

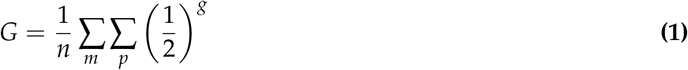

where *n* is the total number of nestlings born that year, *m* is the number of nestlings related to the focal individual, *p* is the number of paths in the pedigree linking the focal individual and the nestling, and *g* is the number of generations separating the focal individual from the nestling in that path (1–3). We used pedigree-based simulations to estimate expected individual genetic contributions instead. Our simulation results match theoretical expectations but also provide estimates of the variance around the expected values.

We used gene dropping simulations to obtain expected genetic contributions of individual breeders and of different immigrant cohorts to our population over time. A founder is by definition any individual in the pedigree whose parents are unknown. Thus, all immigrants are founders. We assigned genotypes to all founders as follows: For individual simulations, we assigned the genotype ‘22’ to the focal individual and ‘11’ to all other founders. To assess the expected genetic contributions of different immigrant cohorts, we assigned immigrants appearing in different years different alleles. We then simulated Mendelian transmission of alleles down the pedigree 1,000,000 times using custom C code. The distribution of allele counts in the nestling cohort each year gives the distribution of expected genetic contributions over time.

### Immigrant contribution projection

The proportion of resident alleles in the birth cohort over time (*r*(*t*)) can be written as:

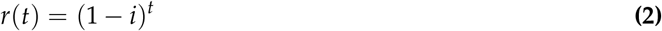

where i is the per-year replacement rate by immigrant alleles and *t* is the number of years following 1992. We began in 1992 because no parents in 1990-1992 were recent immigrants. We fitted this model using non-linear least squares to estimate *i*, then used Eq. (2) to calculate the expected time until neutral alleles are 95% replaced by immigrant alleles.

### Allele frequency predictions

In the absence of selection, the allele frequency of an autosomal SNP in any given year can be written as a function of the individual genetic contributions of each founder and the founder allele frequencies. Let *F* be the number of founders, *G_i,y_* be the expected genetic contribution of founder *i* to the population in year *y*, and *p_i_* be the allele frequency of founder *i*. We can predict the expected allele frequency in year *y* as follows:

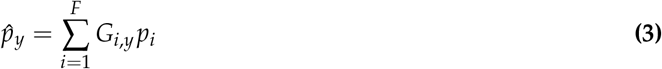

Here we iteratively trimmed the population pedigree until all founders were genotyped and estimated individual genetic contributions using simulations on the trimmed pedigree. We evaluated prediction accuracy by fitting linear regressions.

### Neutral allele dynamics

To generate expected allele frequency distributions over time, we used gene dropping simulations on a trimmed pedigree. For each SNP, we iteratively trimmed the pedigree until all founders in the final trimmed pedigree had a known genotype. Briefly, we removed all ungenotyped founders and set all offspring of these individuals as founders, then repeated these two steps until all remaining founders have observed genotypes. Note that the trimmed pedigree can differ across SNPs because of variable missing data across individuals; however, missing data rates are low (<5%), so these differences are slight. Using the observed genotype for each founder, we simulated Mendelian transmission of alleles down the pedigree a million times and estimated allele frequencies each year in genotyped nestlings from a core set of 54-76 territories.

We used Mann-Kendall tests from the R package Kendall (34) to test for trends in the allele frequencies of incoming immigrant cohorts through time. We tested for net directional selection between 1999-2013 as well as between all adjacent years during that time period by comparing observed allele frequency shifts to the distribution of expected allele frequency shifts generated from the gene dropping simulations. For each test, we calculated *p*-values by counting the number of simulations in which the simulated value is more different from the median value of all the simulations compared to the observed value. We used a FDR threshold of 0.25 for significance.

### Variance in allele frequencies model

To quantify the proportion of variance in the change in allele frequencies due to gene flow and variation in individual survival and reproductive success, we modeled the population as follows: adults who survive or immigrate into the population then produce offspring. From our detailed census and other population monitoring records, we generated a list of individuals present in our population each year in 1990-2013 and categorized them as survivors, immigrants, or nestlings (new births; Fig. S6A). We only included an individual in a given year if it was observed in at least two months during March-June. We conservatively considered individuals who left our study population but later returned as survivors during the intervening time period to minimize inflating the variance in allele frequencies.

Let *N_t_* be the total number of individuals in the population in year *t, N_s_* be the number of individuals who survived from year *t* − 1 to *t, N_i_* be the number of new immigrants into the population in year *t*, and *N_b_* be the number of individuals born in year *t*. Thus the population size in year *t* is *N_t_* = *N_s_* + *N_i_* + *N_b_*. If we denote the allele frequencies in each category as *p_j_*, then we can write the change in allele frequencies between years *t* − 1 and *t* for a given SNP as:

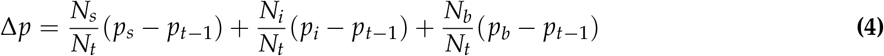

The variance in allele frequency change over time is then:

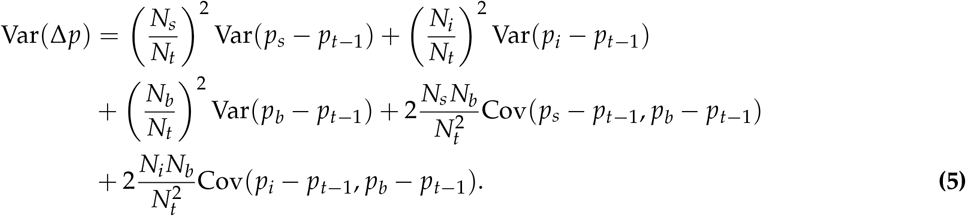

Note that we assume that survivors and immigrants in a given year are unrelated and accordingly set Cov(*p_s_* − *p*_*t*−1_, *p_i_* − *p*_*t*−1_) = 0.

We further partitioned the change in allele frequency due to the birth cohort Var(*p_b_* − *p*_*t*−1_) into the change due to variation in family size and the deviation due to Mendelian segregation of alleles from heterozygotes (Δ*p*_*b*,mend_) (28). If *p_m_* and *p_f_* are the allele frequencies of the parents weighted by the number of offspring they produced in year *t*, then

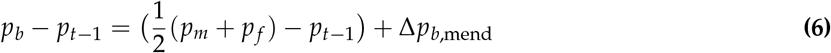

where the first term denotes the expected change in allele frequencies due to the variation in family size and the second the additional independent noise due to Mendelian transmission. We can then estimate the variance due to Mendelian noise as

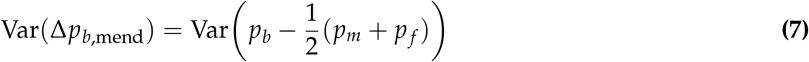

with the alternate term for the variance due to family size variation following from Eq. (6).

We estimated each of the terms on the left and right sides of Eq. (5) averaged across all autosomal SNPs. We then divided each of the terms on the right by the total to quantify the proportion of allele frequency change due to which individuals survive to the focal year, appear as new immigrants, or are born, as well as the contribution of survivors and immigrants to the birth cohort and Mendelian segregation of heterozygotes. We verified our model using simulations.

Though we have genomic data from nearly every individual present in the population from 1999-2013, we still have a small number of ungenotyped individuals in each year (Fig. S6B). To account for missing genotypes, we corrected each term in Eq. (5) for sampling. Normally, the error in allele frequency estimation due to sampling can be statistically modeled, but relatedness among individuals and non-random sampling makes error estimation more complicated in this case. Therefore, we empirically calculated the error in allele frequency estimation using simulations. See SI Text for the full derivation of the model and more details on our simulations. All statistical analyses were done in R (35). All code is available from Figshare.

## Acknowledgments

Thanks to the many students and staff who collected the field data at Archbold Biological Station over the past half century. Thank you Doc Edge for statistical help and the Coopons and Arvid Ågren for comments. This work was supported by NSF (DEB0855879 and DEB1257628) and the Cornell Lab of Ornithology Athena Fund. NC was supported by an NSF Postdoctoral Fellowship in Biology (1523665). NC and GC acknowledge additional support from NIH (R01-GM108779) and NSF (1262327 and 1353380).

## Supplementary Information

### Model derivation

We constructed a model for allele frequency change over time in a population with overlapping generations and fluctuating population sizes. This model relies on the ability to both count all individuals in the population as well as identify new immigrants and new births each year. In our study population, emigration rates are very low, and so we treat emigration events the same as deaths.

Let *N_t_* be the total number of individuals in the population in year *t, N_s_* be the number of individuals who survived from year *t* − 1 to *t, N_i_* be the number of new immigrants into the population in year *t*, and *N_b_* be the number of individuals born in year *t*. Adults who survive or immigrate into the population then produce offspring. Thus the population size in year *t* is *N_t_* = *N_s_* + *N_i_* + *N_b_*.

If we denote the true allele counts in each category *j* as *P_j_* and true allele frequencies as *p_j_* = *P_j_*/2*N_j_*, then the change in allele frequencies (for autosomal SNPs) from year to year is:

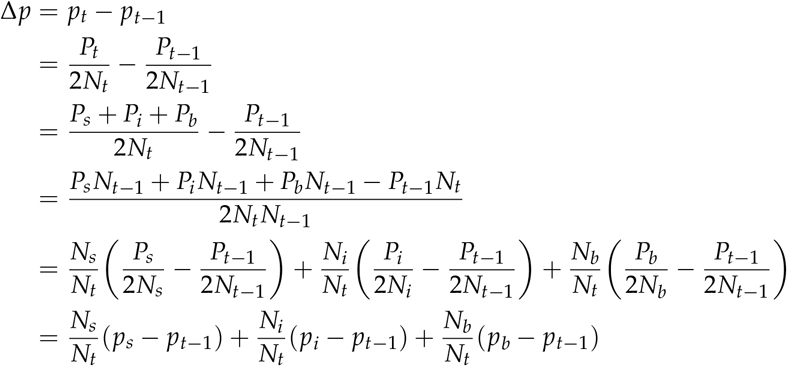

The variance in allele frequency change over time is then:

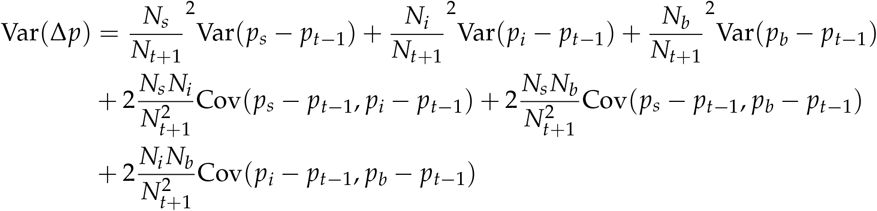

Here, we assume that survivors and immigrants are unrelated despite a few known relationship pairs. We therefore set Cov(*p_s_* − *p*_*t*−1_, *p_i_* − *p*_*t*−1_) = 0. Our model partitions the variance in allele frequency change into contributions from survivors, immigrants, new births, and the covariances between survivors and births as well as immigrants and births:

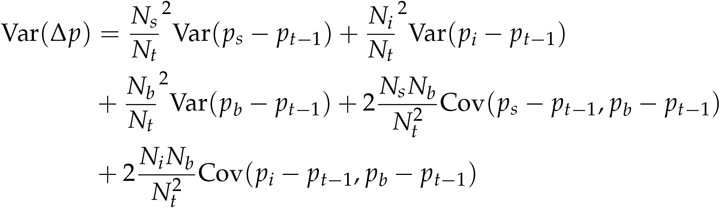

Though we know the numbers of births, deaths, and immigration events each year (*N_j_*), we do not have genomic data from all individuals in the population through time and therefore do not know *p_j_*. We therefore corrected for sampling error as follows. Let *n_j_* be the number of individuals in each category that are genotyped each year, *X_j_* be the observed allele counts, and *x_j_* = *X_j_*/2*n_j_* be the observed allele frequency in each sample. Let *ε_j_* be the error in allele frequency estimation due to sampling (deviation of observed allele frequency from the true unknown allele frequency), such that *x_j_* = *p_j_* + *ε_j_*. Then, using survivors as an example,

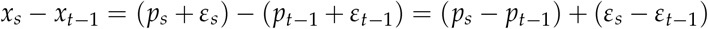

and

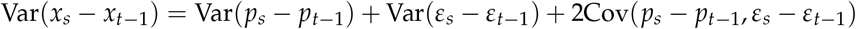

We can solve for Var(*p_s_* − *p_t−1_*):

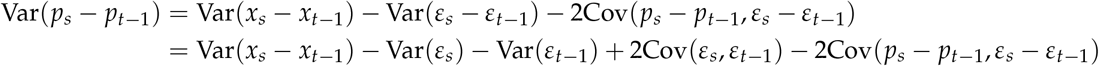

Likewise, the variance terms for immigrants and nestlings are:

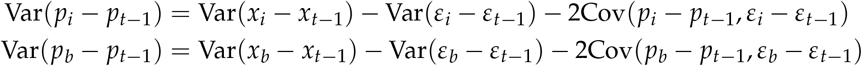

And the two remaining covariance terms are:

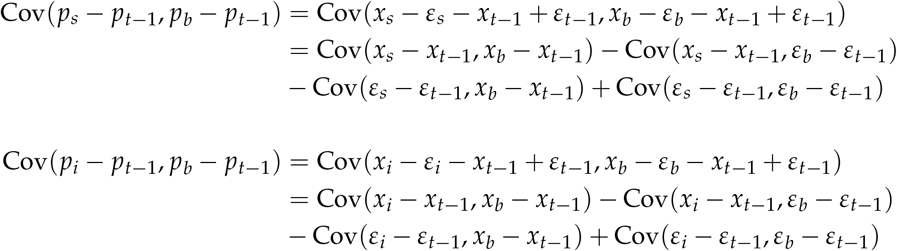

#### Estimation of variance due to Mendelian segregation

The variance in allele frequency change due to births comprises both variation in family sizes and Mendelian segregation of alleles from heterozygous parents. The first term is affected by both genetic drift and selection whereas the second term, Mendelian noise, is a cause of drift. We can partition the proportion of allele frequency change due to these two terms as follows.

In year *t*, the allele frequency of the birth cohort at a given locus is simply the sum of the allele frequency (*g_k_*) of each individual nestling (*k*):

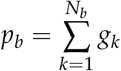

*g_k_* is 0/0.5/1 for individuals with genotype 00/01/11, respectively. From (28), we know that we can write *g_k_* as

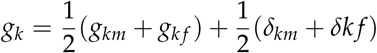

where *g_km_* and *g_kf_* are the allele frequencies of the mother and father of individual *k. δ_km_* and *δ_kf_* is the difference in allele frequency between individual *k* and its parents. The allele frequency of the birth cohort is therefore

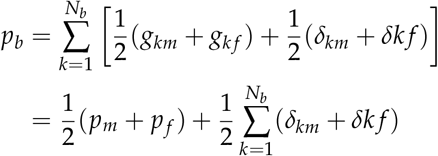

where *p_m_* and *p_f_* are the allele frequencies of all mothers and fathers, respectively, weighted by the number of children they produced in year *t*. For now, let us denote the first term as *p*_*b*,fam_ and the second term as Δ*p*_*b*,mend_. We can estimate the variance in allele frequency change due to variation in family size or segregation in heterozygotes:

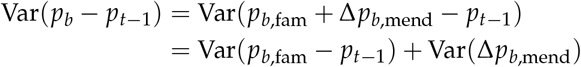

where the covariance is zero by construction. Rearranging Δ*p*_*b*,mend_ in terms of allele frequencies, the variance due to departure from Mendelian transmission of heterozygotes is:

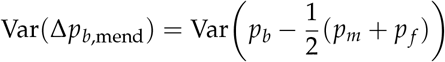

Now, incorporating error due to sampling:

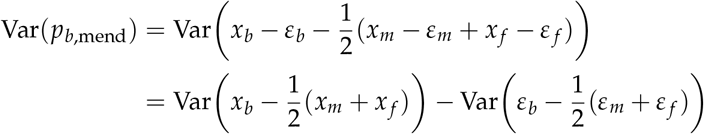

Note that the covariance between the observed estimate and the error is 0. Similarly, the other variance term is:

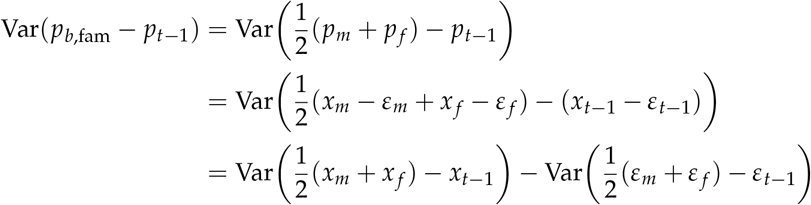

#### Error in allele frequency estimation due to sampling

If sampling is random and individuals are unrelated, then we expect Cov(*ε_s_, ε*_*t*−1_) = Var(*ε*_*t*−1_) (see below for proof) and Cov(*p_s_* − *p*_*t*−1_, *ε_s_* − *ε*_*t*−1_) = 0. The variance in allele frequency change due to survivors then simplifies to:

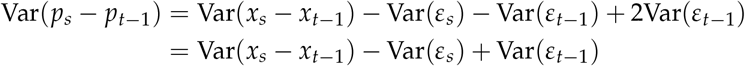

Because *x_j_* is obtained by hypergeometric sampling of 2*n_j_* alleles from a total population of 2*N_j_* alleles with an allele frequency of *p_j_*,

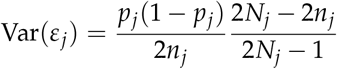

We can estimate heterozygosity from our sample allele frequencies using the small sample size correction 2*n_j_*/(2*n_j_* − 1) from (36).

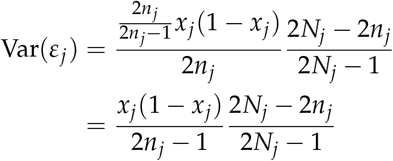

Therefore,

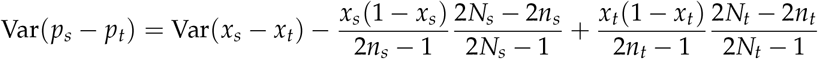

#### Non-random genotyping of parent and children

We noticed that there is non-random sampling in our dataset during the time period of interest (2000–2013). Specifically, immigrants and survivors with at least one offspring are more likely to be genotyped (Fisher’s Exact Test *p* = 6.47 × 10^−5^ and 1.18 × 10^−5^, respectively), and individuals with genotyped offspring are more likely to be genotyped (Fisher’s Exact Test *p* = 0.002 and 1.95 × 10^−6^, respectively). We were more likely to have an archived blood sample from an individual if they had children, particularly in earlier years, before taking blood samples became routine practice in the field. This non-random sampling creates correlations between the error terms for immigrants/survivors and births.

If we define *g_kl_* as a vector of indicator variables denoting whether an individual *k* is genotyped at SNP *l* and *p_kl_* as the allele frequency for individual *k* at SNP *l*, then we can write the sample allele frequency for category *j* as:

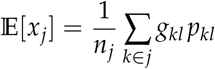

While the true population allele frequency is

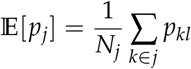

Therefore, the mean error in allele frequency estimation is

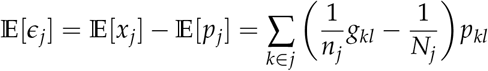

We can write the expected covariance between the error terms for survivors and nestlings as

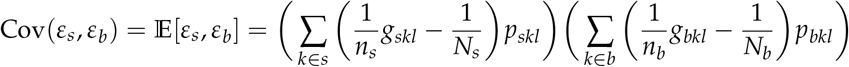

Assuming that non-random sampling occurs only for parent-offspring pairs, then

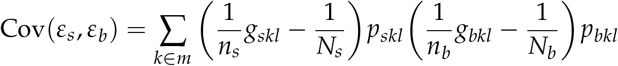

Where *m* is the set of parent-offspring pairs between survivors and nestlings. As the decision to genotype an individual does not depend on their genotype at a specific SNP, then

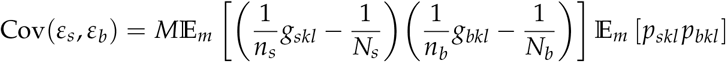

with these expectations being across all *M* parent-offspring pairs.

#### Proof that Cov(ε_s_, ε_t−1_) = Var(ε_t−1_) for random samples

By the law of total covariance,

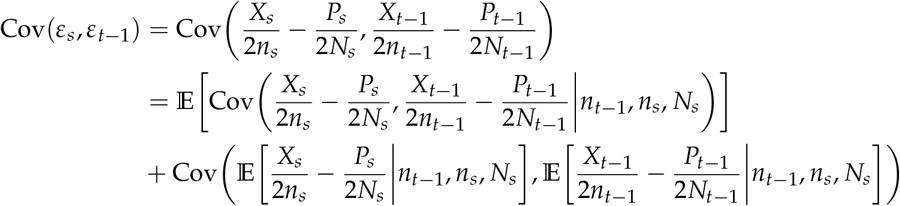

Now 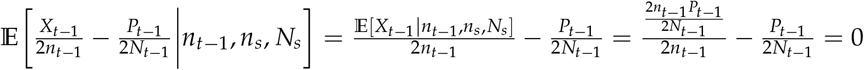, so the second term is 0. As for the first term:

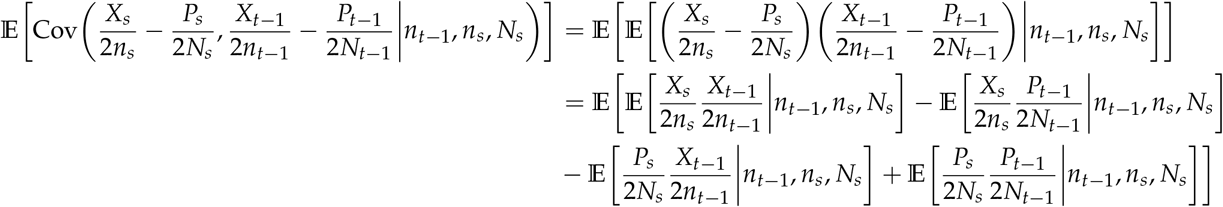

Now we will consider each term separately, dropping the conditions on *n*_*t*−1_, *n_s_, N_s_* for readability.

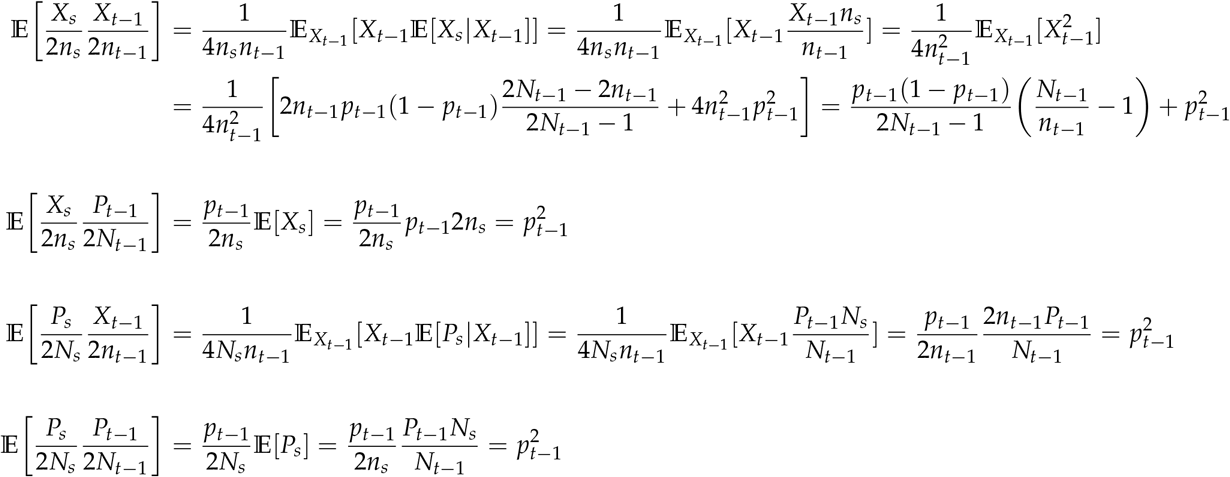

Therefore,

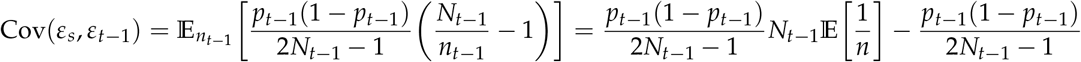

Assuming *n*_*t*−1_ is fixed and not random,

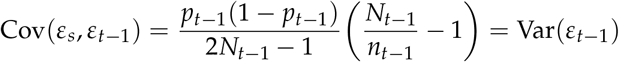

### Implementation

We compiled a list of all individuals present in a core set of territories in 1990-2013. We required a recorded observation in at least two months during March-June to count an individual in a given year. All individuals in the first year, 1990, were considered founders, and individuals were categorized as survivors, immigrants, or nestlings in all subsequent years. Individuals who left our study population but later returned are categorized as survivors during the intervening time period to minimize inflating the variance in allele frequencies. Fig. S6 shows the number of individuals and the proportion who are genotyped in each category for each year.

We then calculated sample allele frequencies for each category in each year at 10,731 autosomal SNPs. Unfortunately, the errors in our case were too complicated to solve analytically, as there was both nonrandom genotyping and relatedness among individuals within and between categories. We therefore used simulations of 100,000 loci using the observed allele frequency spectrum to empirically estimate sampling errors (see below for more details). We averaged across all loci to obtain overall proportions of allele frequency change due to each term. We ran the model for all adjacent years in 1990-2013 but only considered 1999-2013 because we have more genomic sampling during this later time period.

### Simulations

To verify our model, we simulated genotypes for all founders at 10,000 loci using allele frequencies drawn from either a uniform distribution or the observed allele frequency distribution in the first year (1990). We then simulated genotypes for each nestling by randomly drawing an allele from each parent (i.e., simulating Mendelian transmission). Though we simulated genotypes for everyone in the population starting in 1990, we only considered allele frequency changes in the later years with sufficient sampling (same as above). Since we know which individuals are genotyped, we calculated sample allele frequency for each SNP using the subset of genotyped individuals and then subtracted the population allele frequency to get the “true” error in allele frequency estimation. We then ran 1,000 bootstrap iterations, either keeping the sampling scheme constant or changing who is genotyped each time, to verify our model. All analyses were done in the R statistical package (35).

---

**Figure S1.**
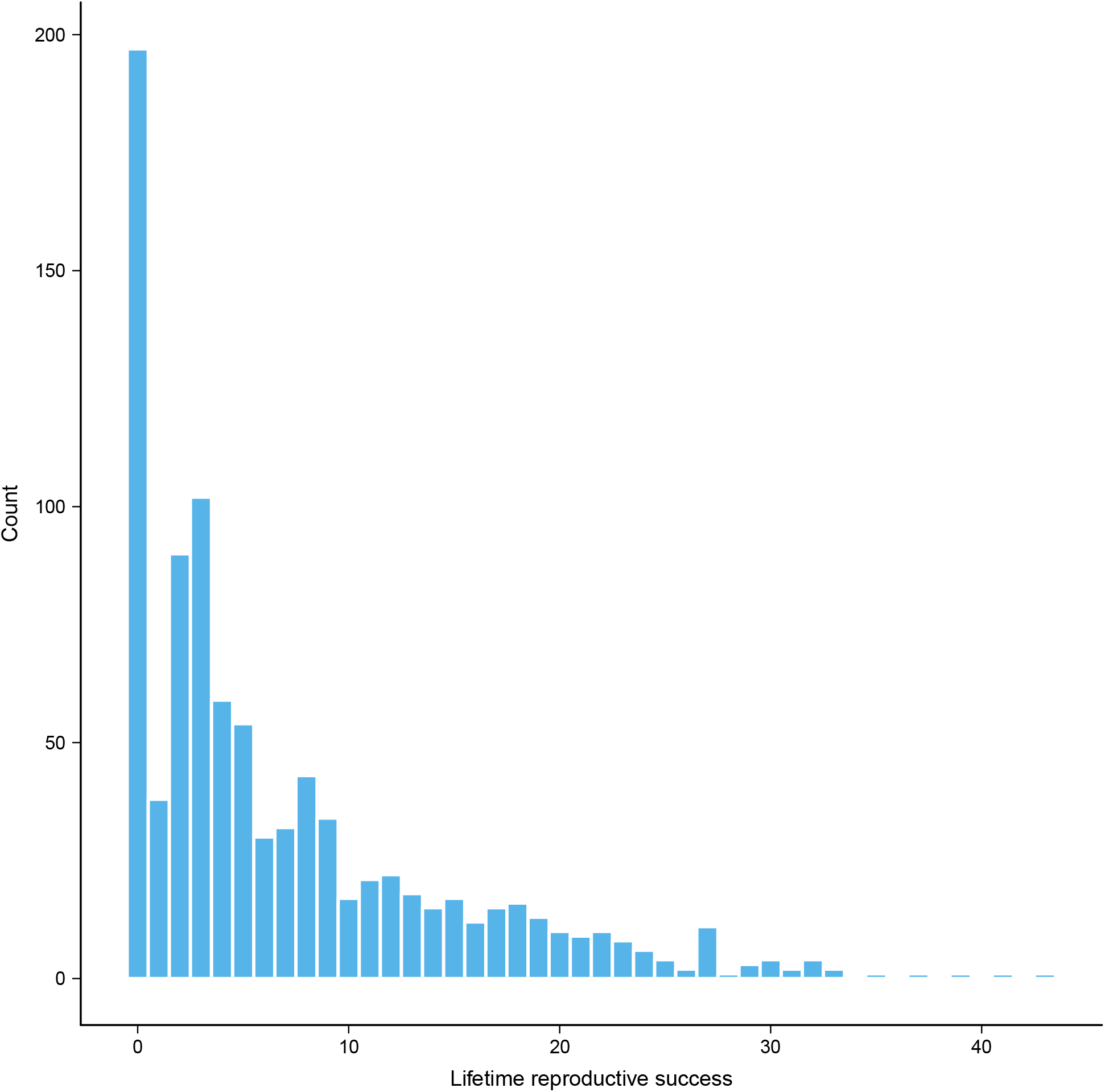
Lifetime reproductive success (measured as the total number of nestlings produced) for all adults born before 2002 and who first bred in 1990 or later (926 individuals).

**Figure S2.**
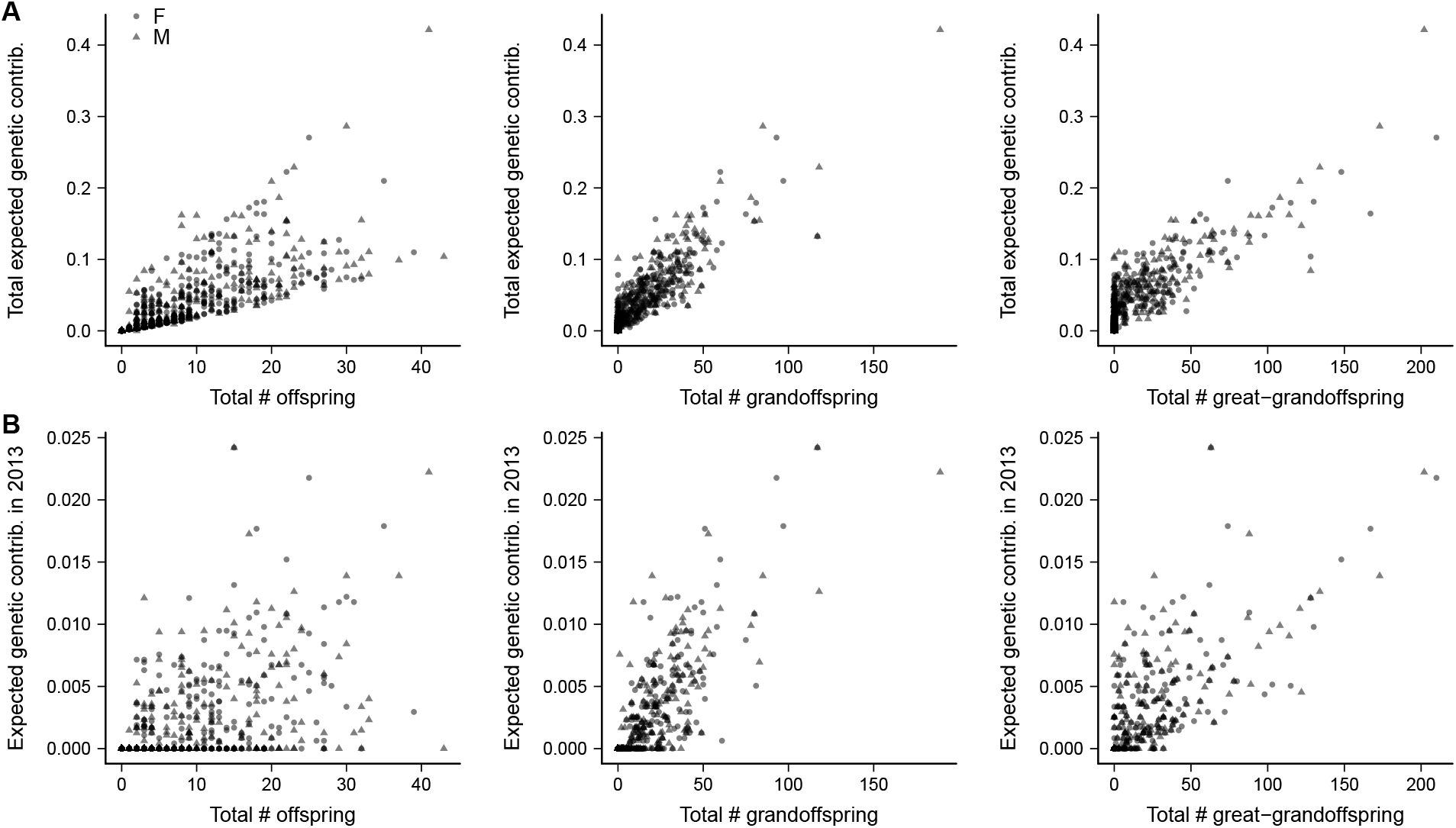
The (A) expected total genetic contribution and (B) expected genetic contribution to the 2013 nestling cohort for an individual is significantly correlated with individual fitness, measured as the total number of offspring, grandoffspring, or great-grandoffspring produced over its lifetime (*p* < 2 × 10^−16^ for all six comparisons). Data are shown for all breeders born before 2002 and who first bred in 1990 or later (926 individuals).

**Figure S3.**
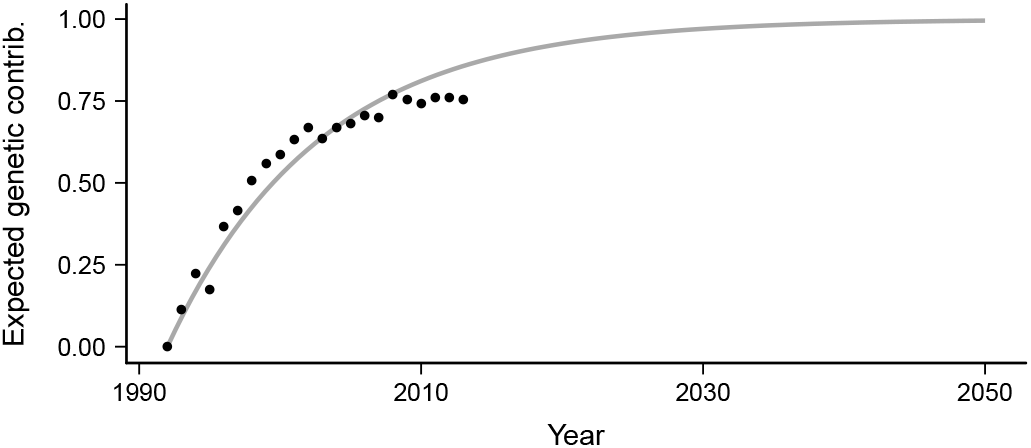
Fitted model for the total expected genetic contribution of immigrants since 1992 over time. Points indicate observed values.

**Figure S4.**
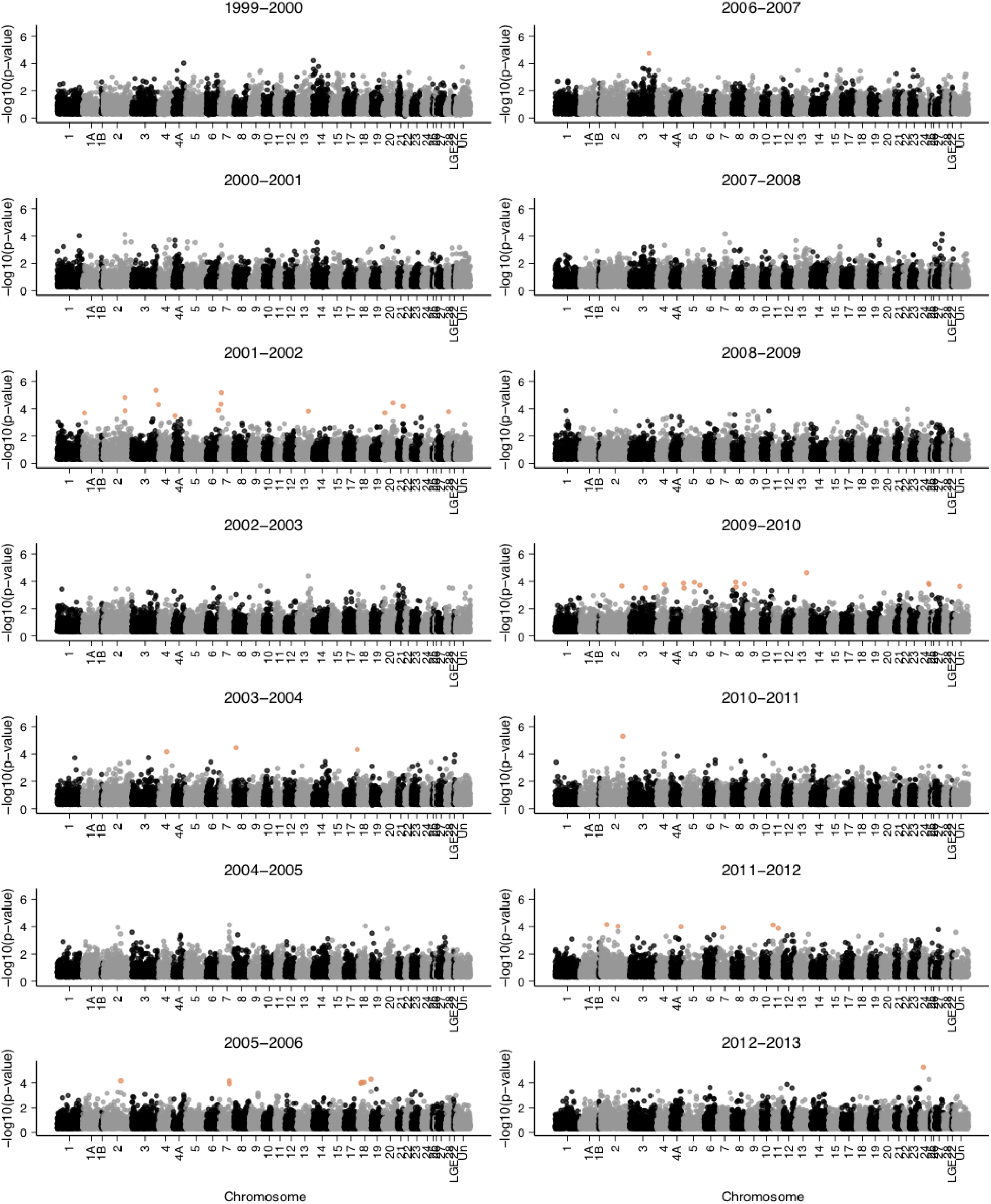
Manhattan plot for allele frequency shifts in all adjacent years from 1999-2013. Significant SNPs (FDR < 0.25) are highlighted in orange.

**Figure S5.**
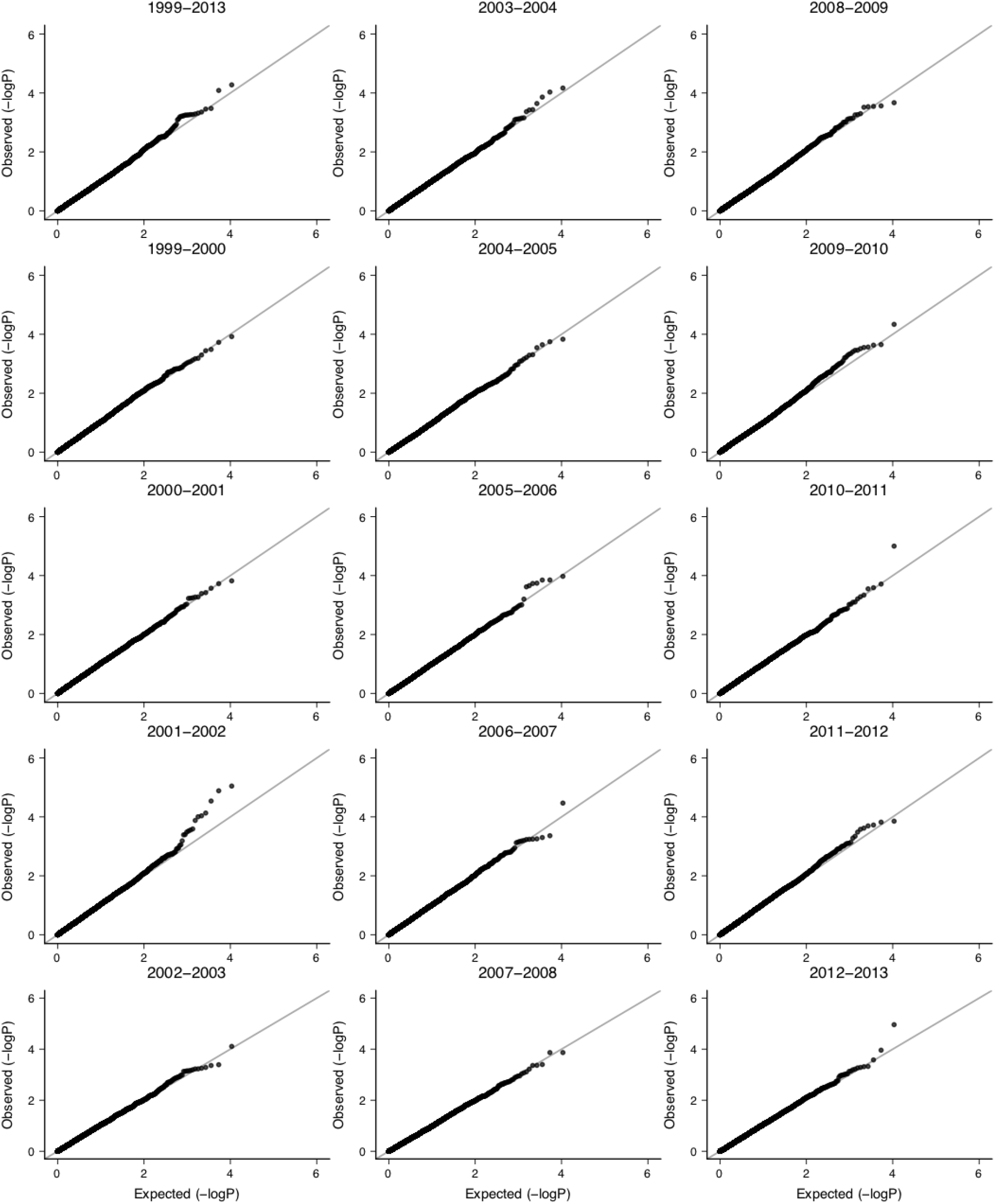
Quantile-quantile plots for all tests of short-term selection.

**Figure S6.**
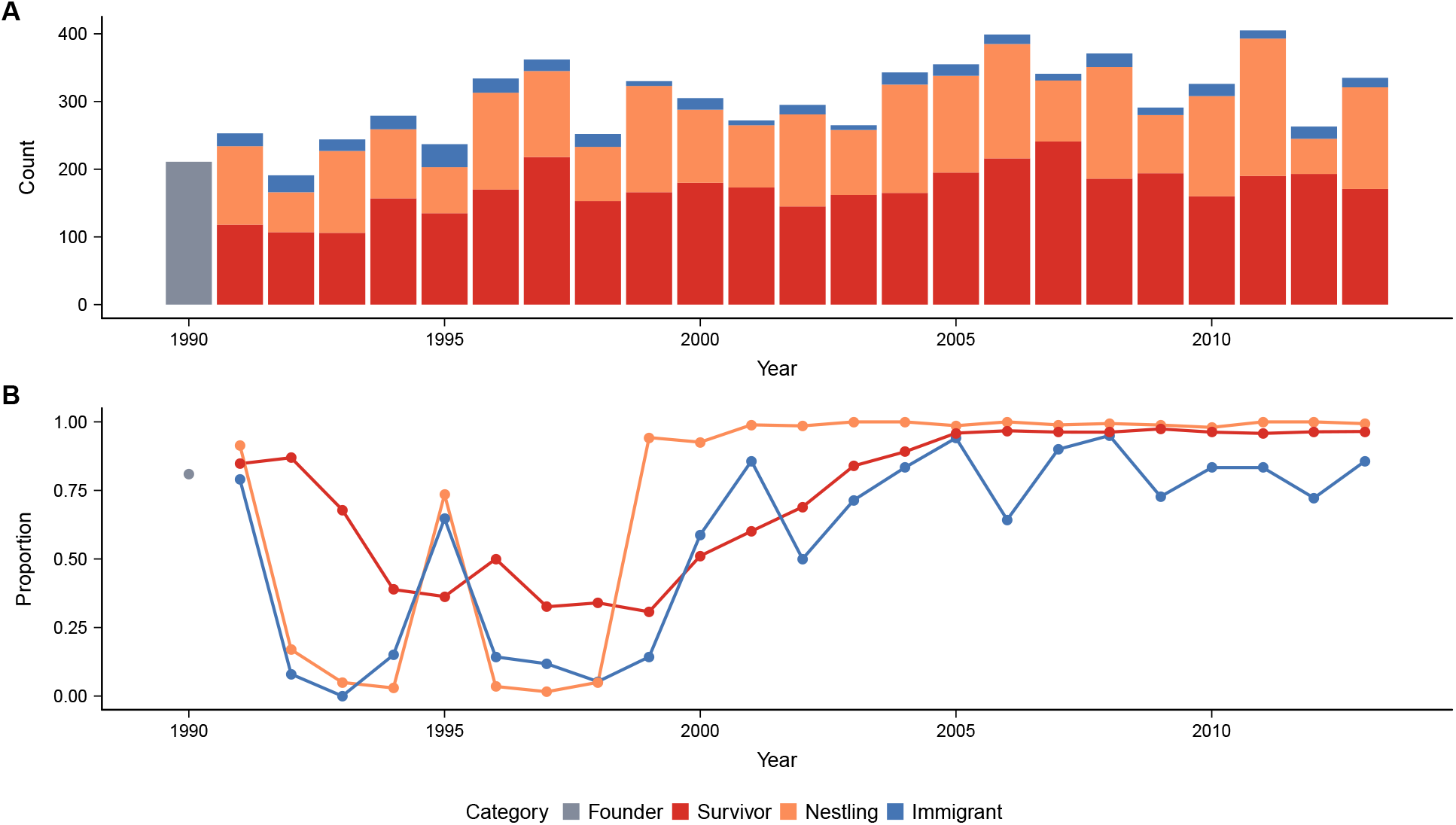
(A) The total number of individuals in each category for the model of allele frequency change over time. (B) The proportion of individuals genotyped in each category each year.

**Table S1.**
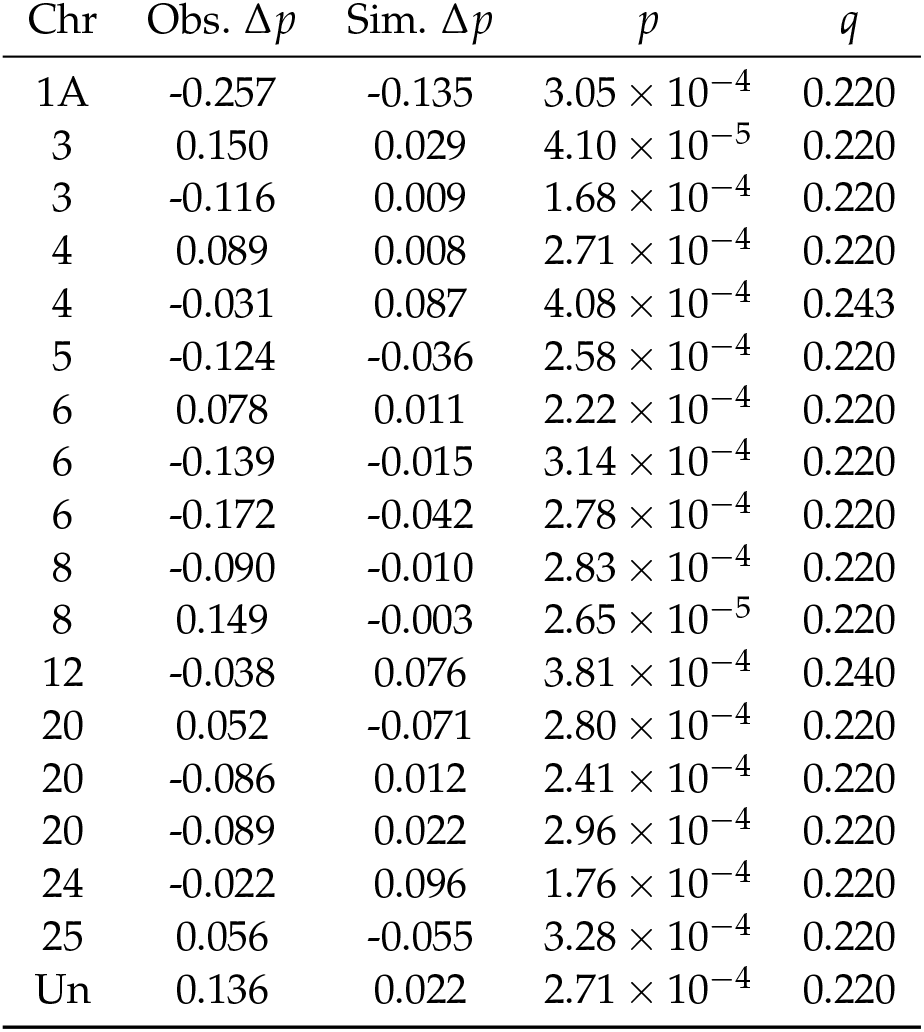
SNPs with significant net allele frequency shifts in 1999-2013.

**Table S2.** SNPs with significant allele frequency shifts between adjacent years.

| Chr | Year | Obs. $\Delta p$ | Sim. $\Delta p$ | $p$ | $q$ |
| --- | --- | --- | --- | --- | --- |
| 1A | 2001-2002 | -0.061 | -0.004 | $2.04 \times 10^{-4}$ | 0.168 |
| 2 | 2001-2002 | -0.065 | 0.014 | $1.45 \times 10^{-5}$ | 0.052 |
| 2 | 2001-2002 | 0.096 | 0.016 | $1.40 \times 10^{-4}$ | 0.160 |
| 3 | 2001-2002 | -0.063 | 0.011 | $4.50 \times 10^{-6}$ | 0.035 |
| 4 | 2001-2002 | 0.034 | -0.003 | $4.95 \times 10^{-5}$ | 0.089 |
| 4A | 2001-2002 | 0.072 | -0.001 | $3.26 \times 10^{-4}$ | 0.249 |
| 6 | 2001-2002 | -0.123 | -0.062 | $1.29 \times 10^{-4}$ | 0.160 |
| 7 | 2001-2002 | 0.051 | -0.044 | $4.60 \times 10^{-5}$ | 0.089 |
| 7 | 2001-2002 | -0.167 | -0.071 | $6.50 \times 10^{-6}$ | 0.035 |
| 13 | 2001-2002 | -0.052 | 0.033 | $1.51 \times 10^{-4}$ | 0.160 |
| 20 | 2001-2002 | -0.060 | 0.036 | $2.00 \times 10^{-4}$ | 0.168 |
| 20 | 2001-2002 | 0.067 | 0.016 | $3.70 \times 10^{-5}$ | 0.089 |
| 21 | 2001-2002 | 0.138 | 0.045 | $6.60 \times 10^{-5}$ | 0.101 |
| 28 | 2001-2002 | 0.026 | -0.040 | $1.64 \times 10^{-4}$ | 0.160 |
| 4 | 2003-2004 | -0.108 | -0.002 | $6.90 \times 10^{-5}$ | 0.247 |
| 8 | 2003-2004 | -0.147 | -0.041 | $3.40 \times 10^{-5}$ | 0.247 |
| 17 | 2003-2004 | 0.074 | -0.031 | $4.65 \times 10^{-5}$ | 0.247 |
| 2 | 2005-2006 | 0.115 | 0.006 | $7.05 \times 10^{-5}$ | 0.185 |
| 7 | 2005-2006 | 0.071 | -0.010 | $7.10 \times 10^{-5}$ | 0.185 |
| 7 | 2005-2006 | 0.065 | -0.004 | $1.21 \times 10^{-4}$ | 0.185 |
| 18 | 2005-2006 | -0.092 | 0.012 | $1.10 \times 10^{-4}$ | 0.185 |
| 18 | 2005-2006 | -0.078 | -0.007 | $9.30 \times 10^{-5}$ | 0.185 |
| 18 | 2005-2006 | 0.129 | 0.016 | $9.05 \times 10^{-5}$ | 0.185 |
| 18 | 2005-2006 | -0.109 | 0.008 | $5.30 \times 10^{-5}$ | 0.185 |
| 3 | 2006-2007 | -0.180 | -0.052 | $1.70 \times 10^{-5}$ | 0.182 |
| 2 | 2009-2010 | -0.104 | -0.012 | $2.23 \times 10^{-4}$ | 0.232 |
| 3 | 2009-2010 | -0.067 | -0.006 | $3.01 \times 10^{-4}$ | 0.239 |
| 4 | 2009-2010 | -0.088 | 0.059 | $1.75 \times 10^{-4}$ | 0.232 |
| 5 | 2009-2010 | -0.073 | -0.021 | $1.36 \times 10^{-4}$ | 0.232 |
| 5 | 2009-2010 | 0.081 | -0.047 | $3.12 \times 10^{-4}$ | 0.239 |
| 5 | 2009-2010 | -0.078 | 0.050 | $1.17 \times 10^{-4}$ | 0.232 |
| 5 | 2009-2010 | -0.132 | 0.016 | $1.96 \times 10^{-4}$ | 0.232 |
| 8 | 2009-2010 | -0.082 | 0.042 | $1.12 \times 10^{-4}$ | 0.232 |
| 8 | 2009-2010 | -0.095 | 0.039 | $2.61 \times 10^{-4}$ | 0.233 |
| 8 | 2009-2010 | -0.071 | -0.016 | $1.54 \times 10^{-4}$ | 0.232 |
| 13 | 2009-2010 | -0.158 | -0.003 | $2.30 \times 10^{-5}$ | 0.232 |
| 24 | 2009-2010 | 0.109 | 0.002 | $1.41 \times 10^{-4}$ | 0.232 |
| 24 | 2009-2010 | 0.058 | -0.031 | $1.74 \times 10^{-4}$ | 0.232 |
| Un | 2009-2010 | -0.117 | -0.002 | $2.38 \times 10^{-4}$ | 0.232 |
| 2 | 2010-2011 | -0.122 | -0.002 | $5.00 \times 10^{-6}$ | 0.054 |
| 2 | 2011-2012 | 0.101 | 0.018 | $7.00 \times 10^{-5}$ | 0.236 |
| 2 | 2011-2012 | 0.167 | 0.010 | $9.45 \times 10^{-5}$ | 0.236 |
| 4A | 2011-2012 | -0.257 | -0.090 | $1.02 \times 10^{-4}$ | 0.236 |
| 7 | 2011-2012 | 0.084 | 0.021 | $1.21 \times 10^{-4}$ | 0.236 |
| 11 | 2011-2012 | -0.198 | -0.020 | $7.55 \times 10^{-5}$ | 0.236 |
| 11 | 2011-2012 | 0.205 | 0.032 | $1.32 \times 10^{-4}$ | 0.236 |
| 24 | 2012-2013 | -0.129 | -0.014 | $5.50 \times 10^{-6}$ | 0.059 |

